# AAV tools enable functional modulation and readout of central and peripheral nervous systems in spiny mice

**DOI:** 10.64898/2026.05.08.723863

**Authors:** Jin Hyung Alex Chung, Renée R. Donahue, Jessica A. Griffiths, Yujie Fan, Changfan Lin, Xinhong Chen, Sayan Dutta, Sarkis K. Mazmanian, Ashley W. Seifert, Viviana Gradinaru

## Abstract

Among mammals, spiny mice (*Acomys* spp.) exhibit the unique capacity to regenerate parts of their nervous system. Studying this phenomenon has the potential to reveal new targets that can slow or halt human neurodegenerative disorders. Unfortunately, research tools (e.g., transgenic lines, gene delivery vehicles) are lacking compared to those available for other rodent models. Here, we tested systemic adeno-associated viral vectors (AAVs) in *Acomys dimidiatus* and identified three promising candidates: X1.1, CAP-Mac, and MaCPNS1. Characterizing their tropism following intravenous delivery, we found that in the brain, MaCPNS1 and X1.1 primarily transduced astrocytes. In the peripheral nervous system, MaCPNS1 efficiently transduced dorsal root ganglia, axon bundles of the ear pinnae, and enteric neurons throughout the gastrointestinal tract. As a proof-of-concept, we used MaCPNS1 to chemogenetically modulate the activity of enteric neurons, successfully decreasing gastric motility in vivo and increasing colonic motility ex vivo. We expect these findings to enable functional studies of the uniquely regenerative nervous system of *Acomys*, which may in turn help advance neuroregenerative therapeutics for humans.

**Summary Statement:** Identification of an AAV tool to efficiently deliver transgenes to the central and peripheral nervous systems of spiny mice enables functional studies of the nervous system in a mammalian model of regeneration.

## Introduction

Spiny mice (*Acomys* spp.) are murid rodents that exhibit enhanced regenerative ability compared to other mammals, a capacity they retain throughout adulthood^1^. For example, they can regenerate skin and musculoskeletal tissue in the ear pinna and exhibit functional regeneration following injury to the kidneys or heart^2–7^. Regeneration is also observed following either hemispinal cord lesion or complete bilateral transection of the spinal cord^8,9^. By contrast, adults of most other mammalian species studied to date, including laboratory mouse (*Mus musculus*) strains and humans (*Homo sapiens*), exhibit limited regenerative ability^10–12^; acute injury is usually repaired with scar tissue, and full functional restoration of the original tissue is rarely achieved^7,9,13–15^. Regenerative healing in mammals is orchestrated by multiple complex processes involving immune-fibroblast crosstalk, nervous inputs, and developmental signaling pathways (e.g., canonical Wnt signaling, ERK activation) working in concert to foster a regenerative wound microenvironment as opposed to one that directs fibrotic repair^8,13,16–19^.

Prior studies comparing regeneration-incompetent mice and regeneration-capable spiny mice have focused on immune responses following acute injuries^8,16,16,18,20,21^. Compared to *Acomys*, the healing environment of *M. musculus* is characterized by significantly higher levels of key pro-inflammatory cytokines (IL-6, CCL2, and CXCL1) during the acute phase following injury. However, inhibition of inflammation during the acute phase via COX-2 inhibition is not sufficient to induce regenerative wound healing in *M. musculus*, suggesting involvement of other factors^16^.

The central nervous system (CNS) and peripheral nervous system (PNS) also play critical roles in shaping the wound environment following an injury. Acute and chronic stress worsen wound healing outcomes in humans and rodents via the hypothalamic-pituitary-adrenal (HPA) axis and autonomic nervous system (ANS)^22–28^. Surgical denervation of peripheral cutaneous nerves impairs ear pinna wound healing in *M. mus*^29^. Furthermore, patients suffering from neurodegenerative disorders such as Parkinson’s disease have an elevated risk of infections in cutaneous wounds, increased incidence of skin-related illnesses, and impaired wound healing^30^. *Acomys* offers promise as a model system to investigate the role of the nervous system in regenerative healing; however, applicable tools to manipulate and monitor the nervous system have been lacking.

Adeno-associated viruses (AAVs) are a potent tool for neuroscience research, offering the ability to deliver up to 4.7 kb of exogenous DNA for expression in target tissues of interest^31^. They are commonly used to deliver genes encoding fluorescent proteins for neuronal morphology studies, chemogenetic and optogenetic tools for neuronal activity modulation, and constructs for gene editing^31–34^. In *Acomys*, direct injection of AAV9, a naturally occurring AAV serotype, has been successfully used to deliver clustered regularly interspaced short palindromic repeats (CRISPR) guides and CRISPR-associated protein 9 (Cas9) for gene editing^31^. Unfortunately, direct injections pose technical and safety hurdles. First, they require invasive surgery, such as craniotomy for delivery to the brain^35–37^. Second, they transduce only a small region directly around the injection site, making them poorly suited for targets that are difficult to access surgically (such as the dorsal root ganglia [DRGs] and sympathetic chain ganglia) or widely distributed (e.g., the enteric nervous system [ENS])^37^. Coverage can be extended by multiple injections^38–40^, but this requires additional surgeries and, if performed sequentially, can induce detrimental immune responses due to neutralizing antibodies^41–43^.

Alternatively, AAVs can be systemically administered via minimally invasive intravenous (i.v.) injection, enabling widespread targeting of tissues. However, systemic delivery of naturally occurring serotypes, such as AAV9, is characterized by low transduction of the CNS, which is protected by the blood-brain barrier (BBB)^44,45^. Our group and others have manipulated the protein sequences on the surface of the AAV capsid to create novel AAV variants with heightened target specificity and efficiency after systemic delivery^45^. Most of this engineering has been done in *M. musculus*, where commercially available transgenic lines have enabled Cre recombinase-dependent strategies that resulted in the development of variants capable of crossing the BBB to target the CNS, such as PHP.eB, CAP-B10, and CAP-B22^46–49^. An RNA-based selection method in non-transgenic *Callithrix jacchus* gave rise to the CNS-targeted variant CAP-Mac^50^. Other engineered variants target the PNS (MaCPNS1 and MaCPNS2)^49,51^.

AAV variants engineered in mice such as X1.1, MaCPNS1, and MaCPNS2 have been shown to transduce cells in the CNS and PNS when delivered systemically to non-human primates^49–51^. However, the tropism of these variants differs greatly between species. Furthermore, variants such as PHP.eB have been shown to exhibit different tropism, with varying degrees of BBB penetrance and transduction of neurons, even across different strains of *M. musculus*^52^, likely reflecting differences in receptor expression patterns between strains and species^53–57^. Thus, the optimal AAV for an application in one species cannot be predicted from performance in others.

Here, we screened seven previously engineered systemic AAVs in *Acomys dimidiatus* and identified one, MaCPNS1, that efficiently transduces the PNS and CNS. We then demonstrated the use of MaCPNS1 for functional chemogenetic modulation of the ENS in an adult, non-transgenic spiny mouse, laying the groundwork for future studies of the nervous system in this key model of mammalian regeneration.

## Results and Discussion

### Engineered systemic AAVs efficiently target the CNS and PNS of spiny mice

To facilitate the development of neural disease models and expand our understanding of central and peripheral nerve regeneration, we sought to identify AAV variants capable of efficiently targeting the CNS and PNS in spiny mice. AAV9, a naturally occurring serotype, was the first AAV identified that can (weakly) penetrate the BBB and transduce a small population of astrocytes and neurons^45^. Subsequent engineering efforts altered the sequence of the AAV9 capsid to boost its efficiency in penetrating the BBB. These efforts created novel variants of AAV9, such as PHP.eB, CAP-B10, and CAP-B22, which efficiently cross the BBB and target the CNS in *M. musculus*^46,48^. However, PHP.eB and CAP-B10 lack the same efficacy in crossing the BBB in other model organisms such as rats and macaques^52,54,55^. More recent engineering efforts have given rise to additional AAV9 variants such as CAP-Mac and X1.1, which show heightened capabilities to penetrate the BBB in non-human primates such as marmosets and macaques^49,50^. MaCPNS1 and MaCPNS2 were engineered from AAV9 to efficiently target the PNS of *M. musculus* and *Macaca mulatta*^51^.

In order to efficiently screen these engineered vectors in spiny mice, we utilized a pooled barcoded capsid approach. In each of the engineered capsids—PHP.eB, CAP-B10, CAP-B22, CAP-Mac, X1.1, MaCPNS1, and MaCPNS2—as well as the parent vector AAV9, we packaged mNeonGreen and a unique DNA barcode under the control of the ubiquitous promoter CAG. The molecular barcode of each vector enabled high-throughput assessment of tissue tropism following systemic administration. We delivered the pooled AAV library via retro-orbital (r.o.) injection to *A. dimidiatus* at 6.25E11 vector genomes (vg) per animal per vector, for a total pooled dose of 5E12 vg per animal, and harvested key tissues 3 weeks post-injection, including the olfactory bulb, forebrain, midbrain, hindbrain, liver, small intestine, and large intestine (n=3, 1-year-old male animals). Viral DNA and transcribed mRNA were extracted, sequenced, and quantified to determine the relative enrichment of each capsid variant across target tissues (Fig. 1A).

**Figure 1.**
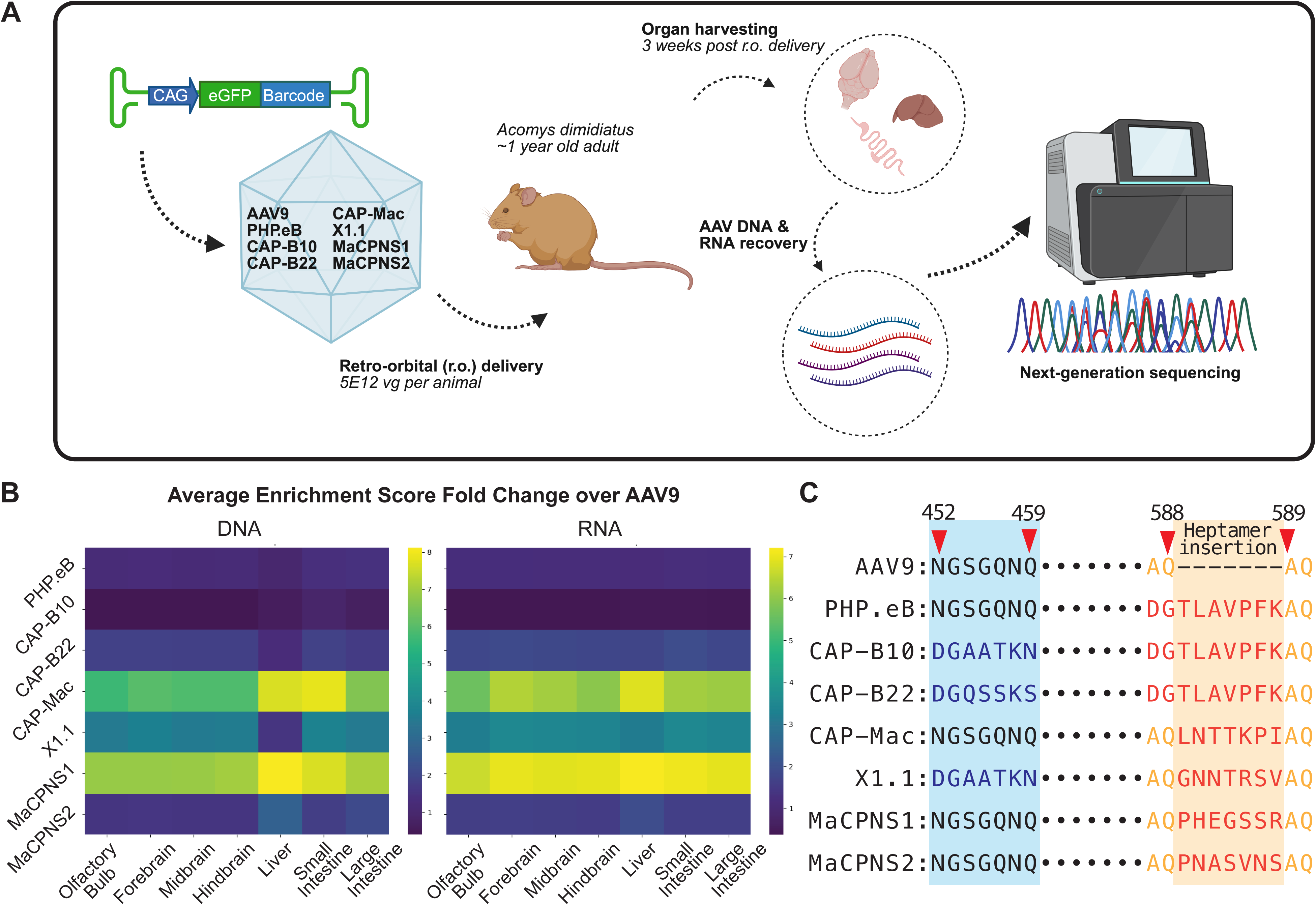
Pooled screening of AAV vectors reveals promising candidates targeting the CNS and PNS of *Acomys dimidiatus* following systemic delivery. **A.** Overview of vector screening. Seven engineered variants of AAV9 (PHP.eB, CAP-B10, CAP-B22, CAP-Mac, AAV9-X1.1, MaCPNS1, and MaCPNS2) along with AAV9, each packaging a uniquely barcoded ssDNA transgene (CAG-mNeonGreen-Barcode) were pooled in equal titers and delivered via retro-orbital (r.o.) injection at a total dose of 5E12 vector genomes (vg) per animal (n=3, 1-year-old males). Three weeks post-delivery, organs of interest were harvested and processed for next-generation sequencing (NGS). **B.** Average enrichment scores of DNA and RNA recovered from various tissues for engineered capsids normalized to scores for AAV9 (n=3, 1-year-old male animals). NGS identified CAP-Mac, X1.1, and MaCPNS1 as promising vectors targeting central (olfactory bulb, forebrain, midbrain, and hindbrain) and peripheral (liver, small intestine, and large intestine) tissues. **C.** Sequences of the engineered regions of variable region (VR)-IV and VR-VIII of the candidate vectors.

Next-generation sequencing (NGS) revealed differential enrichment patterns among the screened capsids (Fig. 1B). Three candidates—CAP-Mac, X1.1, and MaCPNS1—demonstrated high enrichment scores (fold change over AAV9) in tissues of interest, including the CNS and peripheral tissues. Among these, MaCPNS1 showed the most robust and consistent enrichment across multiple regions, including the CNS, small intestine, and large intestine. Interestingly, X1.1, while enriched in several tissue types, exhibited markedly reduced viral DNA levels in the liver relative to CAP-Mac and MaCPNS1, suggesting a difference in tissue-specific transduction efficiency (Fig. 1B). This could potentially be explained by the presence, in the VR-IV loop (amino acids 452–459), of a sequence that weakens the capsid’s interaction with the AAV receptor (AAVR), a receptor hypothesized to be responsible for liver targeting of AAV9-based vectors^55^ (Fig. 1C).

### Systemic AAVs transduce multiple cell types in the CNS of spiny mice

To evaluate the tropism and cellular specificity of the top AAV candidates, CAP-Mac, X1.1, and MaCPNS1, we individually delivered single-stranded (ss) AAV vectors encoding an eGFP reporter under the ubiquitous CAG promoter to *A. dimidiatus*. These vectors, along with the parent AAV9, were administered retro-orbitally at a dose of 2.25E12 vg per animal, and tissue tropism was assessed in sagittal brain sections and peripheral tissues (Fig. 2A, n=3 per vector, 3-month-old female animals).

**Figure 2.**
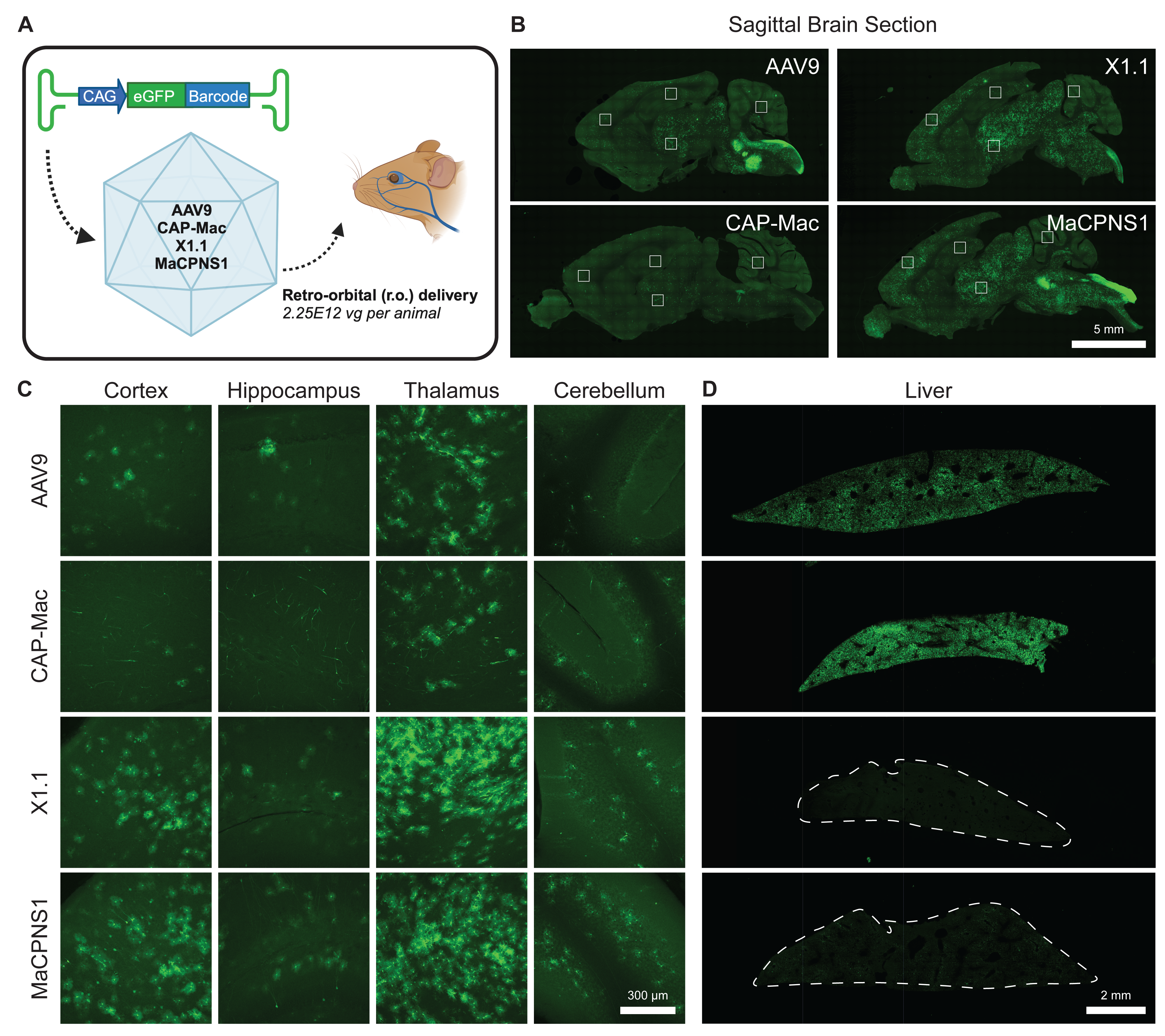
Individual validation of top performing capsids in the CNS and liver. **A.** Schematic of individual vector validation. **B.** Representative images of mid-sagittal brain sections following r.o. delivery of 2.25E12 vg of AAV9, CAP-Mac, X1.1, or MaCPNS1 packaging CAG-eGFP (n=3 per vector, 3-month-old female animals, scale bar = 5 mm). **C.** Enlarged views of boxed areas in **(B)** showing representative regions of cortex, hippocampus, thalamus, and cerebellum (scale bar = 300 μm). **D.** Representative images of livers from animals injected with AAV9, CAP-Mac, X1.1, or MaCPNS1 (scale bar = 2 mm).

Compared to AAV9, both X1.1 and MaCPNS1 displayed markedly enhanced eGFP expression across the brain, suggesting greater transduction efficiency in the CNS (Fig. 2B). CAP-Mac exhibited lower overall eGFP expression in central sagittal sections of the brain compared to AAV9 (Fig. 2B). Both X1.1 and MaCPNS1 produced more abundant eGFP expression in astrocytes within the cortex, hippocampus, thalamus, and cerebellum (Fig. 2C). While MaCPNS1 preferentially transduced astrocytes in the cortex, it also exhibited sparse transduction of neurons (Fig. S1). AAV9 predominantly transduced astrocytes in the cortex and thalamus (Fig. S1).

Astrocytic tropism is advantageous for studies of neuroinflammation, neuron–glia interactions, and regeneration, as astrocytes play pivotal roles in these processes^58–60^. MaCPNS1 could thus serve as a powerful vector for such experiments, especially when combined with an astrocyte-specific promoter such as the glial fibrillary acidic protein (GFAP) promoter^56^. This could enable future work exploring the role of astrocytes in promoting functional recovery following CNS injury in spiny mice.

We also sought to characterize the vectors’ tropism in the liver, where high transduction can induce hepatotoxicity, leading to liver failure and death^62^. CAP-Mac exhibited liver transduction levels comparable to AAV9, as evidenced by widespread eGFP expression (Fig. 2D). However, MaCPNS1 and X1.1 displayed significantly reduced eGFP expression in hepatic tissue, suggesting that they may offer reduced adverse hepatotoxic effects (Fig. 2D).

### MaCPNS1 efficiently and robustly transduces peripheral neurons in Acomys

Building on prior studies demonstrating MaCPNS1’s ability to target the PNS in rodents (*M. musculus* and *Rattus norvegicus*) and non-human primates (*C. jacchus* and *M. mulatta*)^51^, we investigated whether MaCPNS1 exhibits similar PNS tropism in *A. dimidiatus*. To assess this, we examined the transduction of peripheral tissues following systemic delivery of AAV9 or MaCPNS1, focusing on key regions of the PNS.

We first evaluated transduction in the DRGs, a critical hub of sensory neurons. Quantitative analysis of bulk eGFP fluorescence revealed strikingly higher transduction efficiency of MaCPNS1 compared to AAV9 (Fig. 3A–B; mean fluorescence intensity 16.36 a.u. vs. 0.94 a.u.), confirming the superior PNS-targeting capability of MaCPNS1 in *A. dimidiatus*. Further supporting its broad PNS tropism, MaCPNS1 robustly transduced the nerves innervating the ear pinnae in *A. dimidiatus*, producing strong eGFP expression (Fig. 3C).

**Figure 3.**
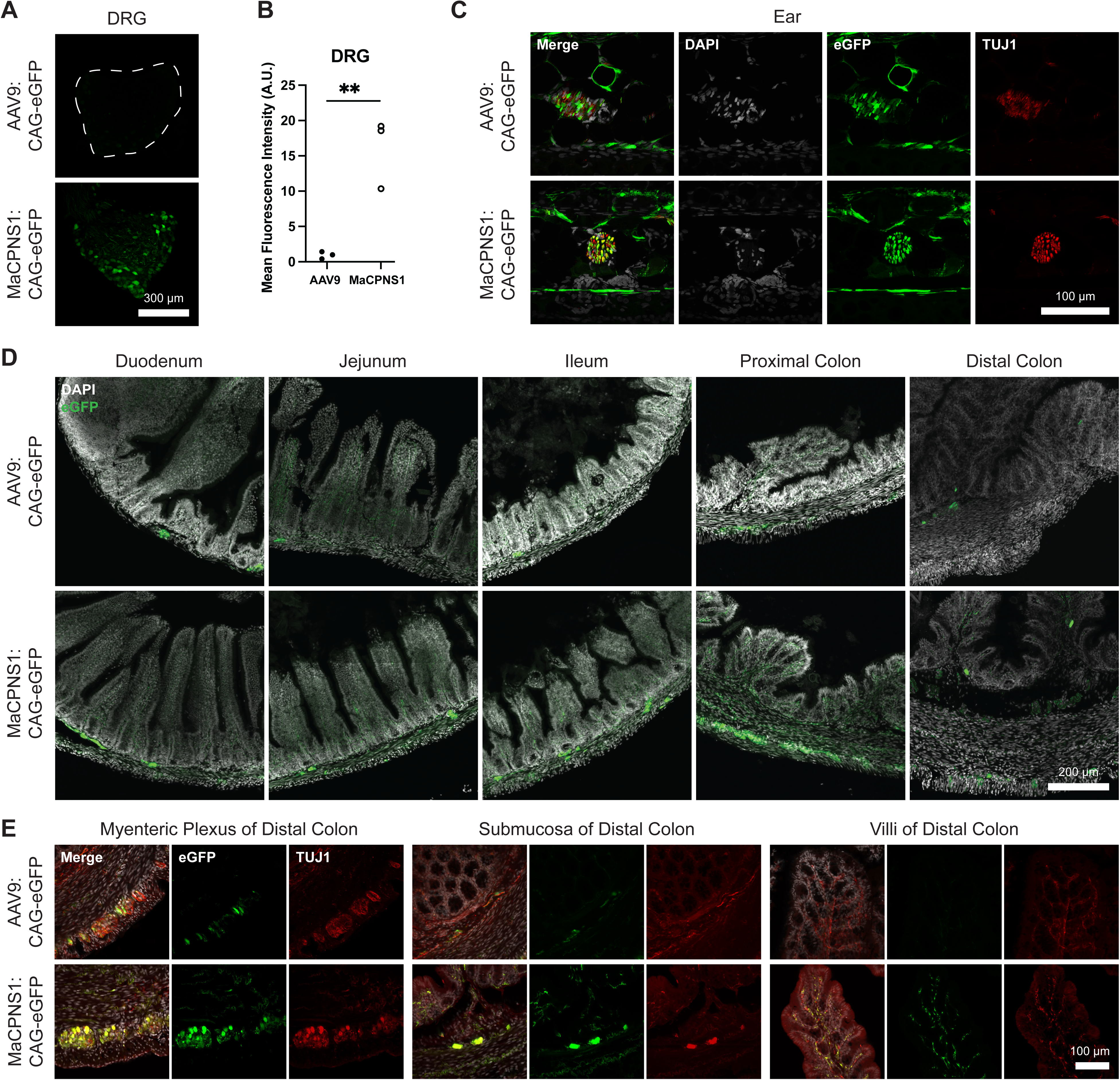
Individual characterization of MaCPNS1 in the PNS. **A.** Dorsal root ganglion (DRG) from thoracic (T2) vertebra following systemic delivery of AAV9:CAG-eGFP (top) or MaCPNS1:CAG-eGFP (bottom) (same animals as in Figure 2, scale bar = 300 μm). **B.** Mean fluorescence intensity in arbitrary units (a.u.) of DRGs of *A. dimidiatus* receiving AAV9 or MaCPNS1 (Student’s t-test, **p < 0.01, n=3 per group, 3-month-old females). Each point represents the measurement from one T2 DRG of one animal. **C.** Cross section of an ear pinna following delivery of AAV9:CAG-eGFP (top) or MaCPNS1:CAG-eGFP (bottom), stained for DAPI (white), eGFP (green), and TUJ1 (red) (scale bar = 100 μm). **D.** Representative images of duodenum, jejunum, ileum, proximal colon, and distal colon of animals injected with AAV9:CAG-eGFP or MaCPNS1:CAG-eGFP, showing eGFP signal (green) and DAPI stain (white) (scale bar = 400 μm). **E.** Representative images of the myenteric plexus, submucosa, and villi of the distal colon of animals injected with AAV9:CAG-eGFP or MaCPNS1:CAG-eGFP showing eGFP fluorescence (green), anti-TUJ1 (red) and DAPI (white in merge) (scale bar = 100 μm).

To examine transduction in the ENS, we analyzed fluorescence in the duodenum, jejunum, ileum, proximal colon, and distal colon. MaCPNS1 demonstrated robust eGFP expression in enteric ganglia and villi throughout all intestinal segments, indicating its capacity to effectively target the ENS in *A. dimidiatus*. In contrast, AAV9 exhibited much lower levels of fluorescence in these tissues (Fig. 3D–E). Confirming MaCPNS1’s tropism for peripheral neurons, the eGFP signal in the distal colon colocalized with the neuronal marker TUJ1 (Fig. 3E).

### MaCPNS1 enables functional modulation of gastrointestinal motility in Acomys

Based on our characterization, we hypothesized that MaCPNS1 could enable functional manipulation of neuronal circuits in *A. dimidiatus*. Previous studies have demonstrated the utility of PNS-targeting AAV vectors for manipulating neuronal subpopulations involved in gastrointestinal (GI) motility in transgenic Cre-expressing lines of *M. musculus*^63,64^. As a proof-of-concept, we therefore explored the potential of MaCPNS1 for functional manipulation of GI motility in adult, non-transgenic *A. dimidiatus* using excitatory designer receptors exclusively activated by designer drugs (DREADDs).

We packaged a broad neuronal promoter (hSyn) driving expression of the excitatory DREADD hM3D(Gq) fused to a fluorescent reporter (eGFP) into MaCPNS1 and delivered the vector into *A. dimidiatus* at a dose of 2.25E12 vg per animal by r.o. injection (3–4-month-old female animals; n=3 animals per cohort for in vivo experiments, n=4 animals per cohort for ex vivo experiments). After 3–4 weeks of expression, we activated the DREADD by administering the agonist compound 21 (C21) intraperitoneally and assessed GI motility (Fig. 4A).

**Figure 4.**
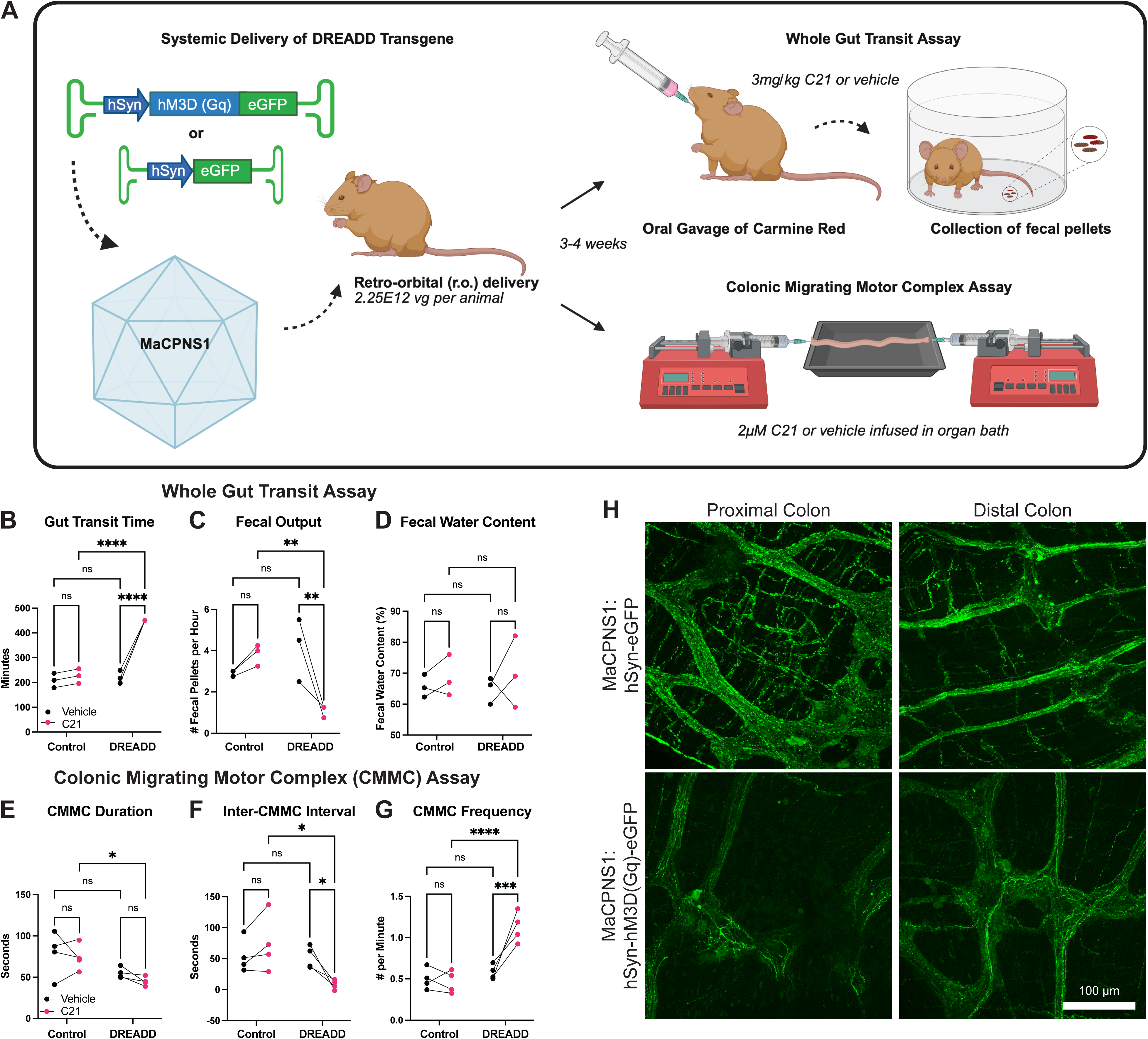
MaCPNS1 efficiently delivers DREADD for functional gut motility modulation. **A.** Schematic of functional gut motility assay utilizing MaCPNS1 packaging a DREADD cargo. Either hSyn-hM3D(Gq)-eGFP or hSyn-eGFP packaged in MaCPNS1 was delivered at 2.25E12 vg per animal via r.o. injection. 3–4 weeks later, on the first day of assessment, animals were administered 300 μL of carmine red followed by i.p. delivery of saline vehicle to measure baseline gut motility. 24 hours later, we administered 300 μL of carmine red followed by i.p. delivery of C21 (agonist for the hM3D receptor, 3 mg/kg, i.p.). On both days, whole gut transit was monitored for 7.5 hours and fecal samples collected for analysis. A separate cohort of animals was treated the same way, sacrificed, and their colons used for an ex vivo colonic migrating motor complex (CMMC) assay. **B-D.** In vivo, activation-induced changes in **(B)** whole gut transit time, **(C)** average fecal pellet output per hour, **(D)** average fecal pellet water content (n=3 3–4-month-old female animals per group). **p < 0.01, ****p < 0.0001, ns not significant, determined by 2-way ANOVA. **E-F.** Ex vivo activation-induced changes in CMMCs measured by **(E)** average duration of individual CMMCs, **(F)** average time between CMMCs, and **(G)** average number of CMMCs per minute (n=4 3–4-month-old female animals per group). *p < 0.05, ***p < 0.001, ****p < 0.0001, ns not significant, determined by 2-way ANOVA. **H.** Representative images of proximal and distal colon stained for eGFP following administration of MaCPNS1:hSyn-eGFP or MaCPNS1:hSyn-hM3D(Gq)-eGFP (scale bar = 100 μm).

Activation of peripheral neurons with C21 significantly reduced gut motility, as evidenced by prolonged whole gut transit time and reduced average fecal output (Fig. 4B–C). Notably, there was no detectable change in the water content of fecal pellets following C21 administration, indicating specific modulation of motility rather than water reabsorption (Fig. 4D).

We also assessed colonic migrating motor complexes (CMMCs) ex vivo using excised colon tissue in an organ bath. While C21 administration did not alter the duration of individual CMMCs, it markedly decreased the inter-CMMC interval and increased the frequency of CMMCs per minute (Fig. 4E–G). These findings suggest that MaCPNS1-delivered DREADD can modulate colonic motility, likely through enhanced neuronal excitability, and highlight the potential of MaCPNS1 for delivering functional cargos to the PNS to investigate various physiological systems influenced by peripheral nerves.

The apparent discrepancy between the prolonged whole gut transit time in vivo and increased frequencies of CMMCs ex vivo can potentially be explained by different distributions of various neuronal subpopulations throughout the gastrointestinal tract, as has been observed in *M. musculus*^58^. Activation of choline acetyltransferase-positive (ChAT+) enteric neurons in *M. musculus* increases both fecal output and CMMC frequency^63^. However, activation of tyrosine hydroxylase-positive (TH+) enteric neurons increases fecal output without changing CMMC frequency^63^. Nitric oxide synthase-positive (NOS+) neurons of the ENS facilitate smooth muscle relaxation and dysfunction of NOS+ neurons can lead to gastroparesis in mice^65^. The proper orchestration of the activity of TH+, ChAT+, and NOS+ neurons is crucial for proper gut motility. Here, by non-specifically activating PNS neurons, we may have disrupted this balance, leading to the disruptions in gut motility we observed. Future studies employing subtype-specific promoters and enhancers in combination with MaCPNS1 may uncover the functional roles of different subpopulations of enteric neurons in *Acomys*.

The eGFP reporter conjugated to the transgene showed robust fluorescence in the proximal and distal colon, confirming widespread expression of the DREADD in the PNS following systemic delivery (Fig. 4H). Since this experiment used the neuronal hSyn promoter, we also examined eGFP expression in the CNS, to compare it to the expression pattern observed earlier with the CAG promoter. Sparse expression was evident in neuronal populations across the brain, confirming that MaCPNS1 does transduce some neurons in the CNS of *A. dimidiatus* (Fig. S2).

Together, this study significantly advances the genetic toolkit available for *A. dimidiatus*, demonstrating that AAV-MaCPNS1 can achieve efficient, systemic delivery of functional cargos to the PNS and CNS of spiny mice. The ability of MaCPNS1-delivered DREADD to functionally modulate GI motility demonstrates the potential of this vector for studying neural circuits and peripheral nerve function. As a non-invasive and versatile tool, MaCPNS1 lays the foundation for expanding the use of *Acomys* as a model system in regenerative biology and beyond.

## Materials and Methods

### Animals

All animal procedures carried out in this study were approved by the California Institute of Technology Institutional Animal Care and Use Committee (Caltech IACUC), Caltech Office of Laboratory Animal Resources (OLAR), and the University of Kentucky IACUC. Male and female spiny mice (*Acomys dimidiatus*), 2–4 months old, were obtained from in-house breeding colonies at Caltech and the University of Kentucky. Animals were housed with ad libitum access to food and water under a 12:12-hour light/dark cycle. For all experiments performed in this study, the animals were randomly assigned to groups.

### AAV Vector Design and Production

AAV packaging and purification were carried out as previously described, with modifications to utilize AAV-MAX suspension cells (Thermo Fisher, A51217, dx.doi.org/10.17504/protocols.io.kqdg31kzzl25/v1)^66^. In brief, recombinant AAV was produced by triple transfection of suspension cells using the VirusGEN AAV Kit with RevIT enhancer (Mirus Bio, MIR 8007) at a molar ratio of transgene: AAV capsid: pHelper = 1:2:0.5, as per the manufacturer’s instructions. The total DNA amount was 2 μg per mL of cells. Virus-producing cells and medium were harvested 72 hours post-transfection. Viral particles were purified using iodixanol step gradient columns followed by ultracentrifugation, as previously detailed^66^. The purified AAV was quantified through droplet digital PCR (Bio-Rad) following Addgene’s protocol with modifications (dx.doi.org/10.17504/protocols.io.261gery8dl47/v1). Briefly, AAV samples were serially diluted in nuclease-free water to achieve a suitable concentration for ddPCR analysis. The reaction mixture was prepared using 10 μL of ddPCR Supermix for Probes (Bio-Rad), 1 μL of primers binding to ITR regions (final concentration 900 nM), 5 μL of the diluted AAV sample, 10 μL of 2X ddPCR EvaGreen supermix (Bio-Rad) and nuclease-free water to a final volume of 20 μL. The mixture was then partitioned into droplets using the QX200 Droplet Generator (Bio-Rad). Thermal cycling was performed under the following conditions: 95°C for 10 minutes, followed by 40 cycles of 95°C for 30 seconds and 60°C for 1 minute, and a final hold at 90°C for 5 minutes. After cycling, droplets were read using a QX200 Droplet Reader (Bio-Rad), and the concentration of positive droplets was analyzed using QuantaSoft software (Bio-Rad). The viral genome titer was calculated using the Poisson distribution, accounting for dilution factors and sample volumes.

Barcode Library Preparation: For pooled screening, a library of barcoded transgenes was designed to facilitate tissue-specific analysis of capsid tropism. Each AAV vector carried a unique 20-nucleotide barcode (BC) downstream of the ubiquitous CAG promoter and mNeonGreen reporter (CAG-mNeonGreen-BC).

Reporter Constructs: For individual characterization, AAV genomes encoded the eGFP reporter driven by the CAG promoter (CAG-eGFP) or the neuronal-specific human synapsin (hSyn) promoter (hSyn-eGFP).

DREADD Constructs: Designer receptors exclusively activated by designer drugs (DREADD; hM3Dq) were fused to eGFP and driven by the hSyn promoter to enable functional studies of peripheral nervous system (PNS) activity (hSyn-hM3Dq-eGFP).

### Systemic AAV Administration

For systemic AAV administration, animals were anesthetized with isoflurane and monitored during and after injections to ensure welfare (dx.doi.org/10.17504/protocols.io.36wgqnw73gk5/v1). Pooled AAV vectors were administered via the retro-orbital sinus at a dose of 5E12 vector genomes (vg) per animal. For pooled experiments, the library was co-injected as a single mixture with equal titers of individual variants. Individual AAV vectors were administered via the retro-orbital sinus at a dose of 2.25E12 vector genomes (vg) per animal.

### Tissue Harvesting and Processing

Four weeks post-injection, animals were anesthetized with isoflurane and transcardially perfused with phosphate-buffered saline (PBS) followed by 4% paraformaldehyde (PFA) for tissue fixation. Dissected tissues included the brain (olfactory bulb, forebrain, midbrain, hindbrain), dorsal root ganglia (DRGs), ear pinnae, liver, small intestine, and large intestine.

Bulk Analysis: For pooled experiments, tissues were homogenized, and DNA and RNA were extracted using the Qiagen AllPrep DNA/RNA Kit. Barcoded AAV sequences were amplified by PCR for DNA or reverse transcription-PCR (RT-PCR) for RNA and sequenced using an Illumina platform. Sequences can be found in the following repository: https://doi.org/10.5281/zenodo.17247291. Sequences were aligned and analyzed using custom scripts generated with Python 3 (https://doi.org/10.5281/zenodo.17247271).

Enrichment scores were calculated using the following equation:

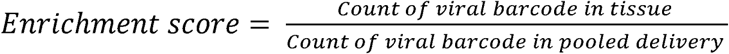

We calculated the fold change of the enrichment score of engineered capsids (PHP.eB, CAP-B10, CAP-B22, CAP-Mac, X1.1, MaCPNS1, and MaCPNS2) in each tissue from the enrichment score of AAV9 using the following equation:

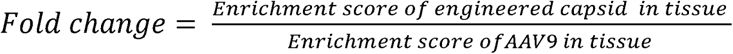

Immunohistochemistry: For all immunohistochemistry (IHC), we followed a modified version of the following protocol (dx.doi.org/10.17504/protocols.io.5qpvokmq7l4o/v1) appropriate for each of the following tissue types.

The GI tract was post-fixed in 4% PFA overnight at 4 °C and stored in PBS + 0.025% sodium azide. DRGs were cryoprotected in 10% and then 30% sucrose in PBS for 1 day each. Tissues were embedded and flash frozen in OCT and cryosectioned into 40 μm sections.

Brain tissues were vibratome sectioned into 100 μm sections. After sectioning, the brain sections were stored in PBS + 0.025% sodium azide.

For IHC, sections were blocked in 5% normal donkey serum (Jackson ImmunoResearch Labs Cat# 017-000-121, RRID:AB_2337258) with 0.1% Triton X-100 (Sigma-Aldrich, 93443). Primary and secondary antibodies were diluted in this blocking buffer. Tissue was incubated with primary antibody overnight at 4 °C and with secondary antibody for 2 h at room temperature. After each antibody incubation step, sections were washed three times for 10 min each in 1× PBS with 0.1% Triton X-100. For DAPI labeling, sections were incubated for 15 min with 1/10,000 DAPI 33342 (Thermo Fisher Scientific, 62248) following secondary antibody incubation in 1× PBS, followed by three washes in 1× PBS.

The following primary antibodies and dilutions were used: rabbit anti-NeuN (1:500; Abcam Cat# ab177487, RRID:AB_2532109), rabbit anti-Glut1 (1:500; Millipore Cat# 07-1401, RRID:AB_1587074). Alexa Fluor 594 conjugated goat anti-rabbit secondary antibody (Abcam Cat# ab150080, RRID:AB_2650602) was used at a 1:1,000 dilution.

Tissues were then transferred to slides and coverslipped with Prolong Gold (Molecular Probes P36970). Images were acquired on a Zeiss LSM 780 microscope, and laser settings, contrast, and gamma were kept constant across images that were directly compared. All confocal images were taken with the following objectives: M27 10× objective and Plan-Apochromat 25× objective with oil immersion.

Ear pinnae were post-fixed in 4% PFA, washed with PBS, and then cryoprotected in 30% sucrose in PBS until equilibrated. Tissues were embedded in OCT and flash frozen in isopentane in a liquid nitrogen bath. Pinnae were sectioned at 40 μm on a cryostat and placed into 1 × PBS. Free-floating immunofluorescence was performed for smooth muscle actin (α-SMA, 1:400 dilution, Novus Cat# NB300-978, RRID:AB_2273628) and tubulin β3 (α-TUJ1, 1:200 dilution, BioLegend Cat# 801202, RRID:AB_2313773), followed by secondary antibodies, donkey anti-goat Alexa Fluor 647 (1:1,000 dilution, Invitrogen, now Thermo Fisher Scientific Cat# A-21447, RRID:AB_2535864) and donkey anti-mouse Alexa Fluor 568 (1:1,000 dilution, Invitrogen, now Thermo Fisher Scientific Cat# A10037, RRID:AB_11180865), respectively. Tissues were then transferred to slides, coverslipped with Prolong Gold with DAPI (Molecular Probes P36931) and imaged on a Nikon AR1 Confocal with a 40x objective, using Nikon Elements Software.

Expression of DREADDs was confirmed by staining with rabbit anti-GFP antibody (Invitrogen, A-31852) at a dilution of 1:1,000 in sagittal brain sections and whole-mounted distal colon.

Fluorescence Quantification: Bulk fluorescence intensity in the DRGs and other tissues was quantified using ImageJ. Mean fluorescence intensity was calculated from three independent animals per group.

### Gastrointestinal (GI) Motility Assays

Procedures described previously^63^ were adapted for use in spiny mice.

Whole Gut Transit (dx.doi.org/10.17504/protocols.io.36wgq3p1xlk5/v1): 6% (w/v) carmine red (Sigma-Aldrich, St. Louis, MO) with 0.5% methylcellulose (Sigma-Aldrich) was dissolved and autoclaved prior to use. To assess GI motility, animals receiving DREADD under the hSyn promoter were evaluated 3–4 weeks post-injection. Spiny mice were administered C21 (3 mg/kg) or vehicle (0.9% NaCl in H_2_O) intraperitoneally, and subsequently orally gavaged with 300 μL of carmine red solution. Spiny mice were single housed with no bedding for the duration of the experiment, and animals were not fasted beforehand. Over the 7.5 h following gavage, the time of expulsion was recorded for each fecal pellet, and each pellet was collected in a pre-weighed, 1.5 mL microcentrifuge tube. Each pellet collected was checked for the presence of carmine red, and the time of initial carmine red pellet expulsion was recorded as the whole gut transit time. The mass of collected fecal pellets was determined, and pellets were left to dry in an 80 °C oven for 24 hours before weighing the desiccated pellets and calculating the pellets’ initial water content. Fecal output rate for each mouse was calculated as the total number of pellets expelled during the 7.5 h time course post-C21 administration divided by the time (in hours) when the last fecal pellet was expelled.

Ex vivo Colonic Motility (dx.doi.org/10.17504/protocols.io.n92ldm61nl5b/v1): Intact colons were dissected from cervically dislocated spiny mice, flushed and placed in pre-oxygenated (95% O_2_, 5% CO_2_) Krebs-Henseleit solution at RT. Proximal and distal ends were cannulated to 2 mm diameter tubes and secured in the center of an organ bath with continuously oxygenated Krebs-Henseleit solution at 37 °C. Syringe pumps were connected to the inlet and outlet tubes to maintain a flow of solution at a rate of 0.5 mL/min through the colon. The system was allowed to equilibrate for 30 min before recording. Baseline recordings were taken for 30 min, then the Krebs solution in the organ bath was briefly removed, mixed with C21 to a final concentration of 2 mM, and replaced in the organ bath. Recordings were taken for another 30 min. The number of CMMCs per minute was calculated before and after C21 administration, as well as the duration of individual contractions and the inter-CMMC intervals.

### Data Analysis

All quantitative data were analyzed using GraphPad Prism. Statistical comparisons were performed using unpaired t-tests or two-way ANOVA. Statistical significance was set at *p* < 0.05.

## Acknowledgements

We thank the entire Gradinaru laboratory for careful review and helpful discussions, especially C. Oikonomou for thorough review of the manuscript. We thank P. Anguiano, E. Mackey, and Z. Qu for administrative assistance. We thank T.F. Miles, M. Zhang, J. Hoang, G. Coughlin, and C. Arokiaraj for insightful discussions and guidance on in vivo screening of AAV vectors. This research was funded in whole by Aligning Science Across Parkinson’s (ASAP-020495 to V.G.) through the Michael J. Fox Foundation for Parkinson’s Research (MJFF). For the purpose of open access, the authors have applied a CC BY public copyright license to all Author Accepted Manuscripts arising from this submission.

## Data and Resource Availability

The data, code, protocols, and key lab materials used and generated in this study are listed in a Key Resource Table alongside their persistent identifiers in Supplementary Table 1. All data cleaning, preprocessing, analysis, and visualization were performed using Python 3, Microsoft Excel and GraphPad Prism.

---

**Supplementary Figure 1:**
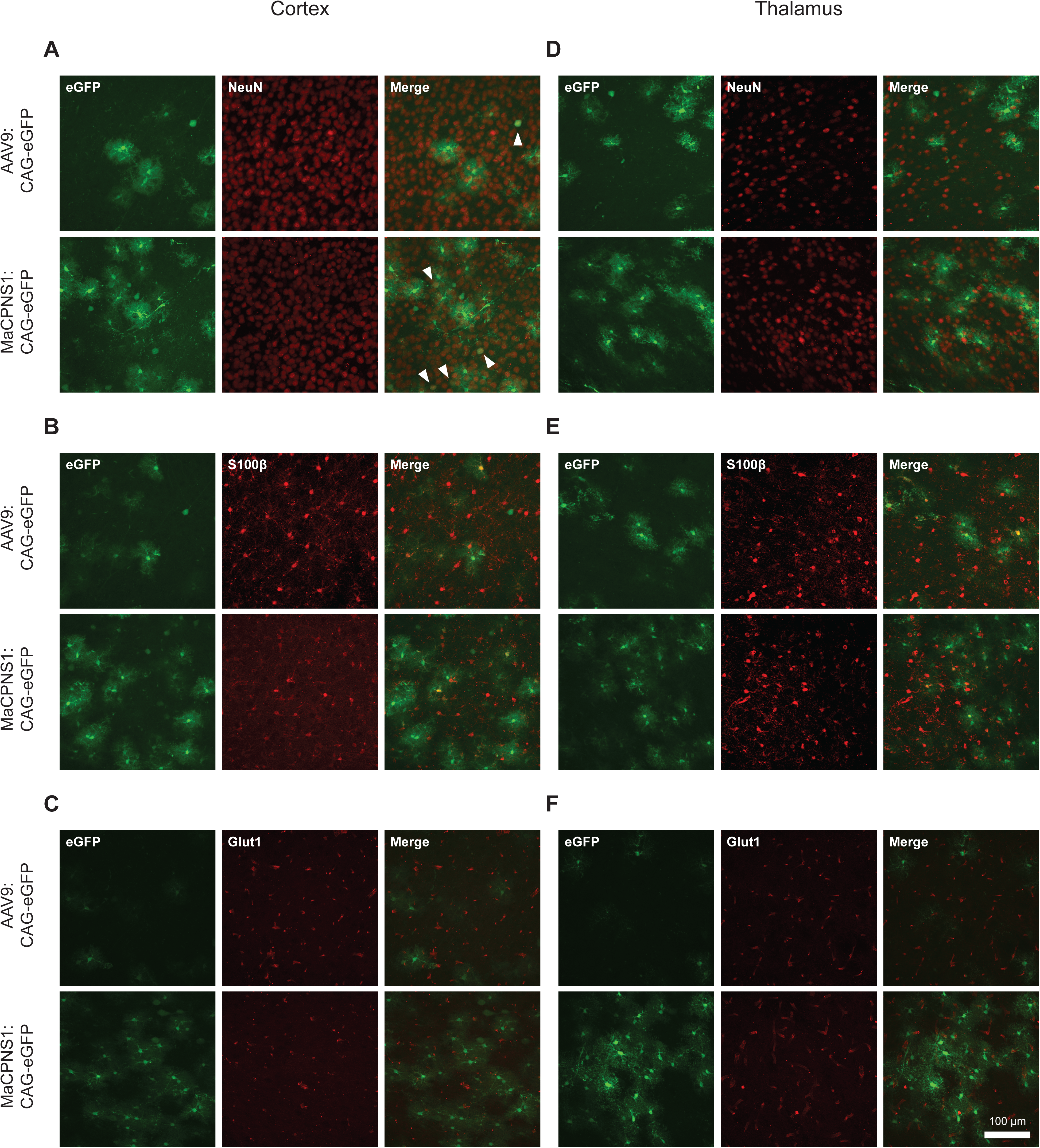
Characterization of AAV9:CAG-eGFP and MaCPNS1:CAG-eGFP transduction in the *A. dimidiatus* brain. Representative images of the cortex **(A-C)** and thalamus **(D-F)** of the same animals as in Figure 2 systemically injected with either AAV9:CAG-eGFP (top) or MaCPNS1:CAG-eGFP (bottom), stained with anti-NeuN (**A, D)**, anti-S100β **(B, E)**, or anti-Glut1 **(C, F)**. White arrows point to eGFP+ co-labeled with anti-NeuN **(A)**. Scale bar = 100 μm.

**Supplementary Figure 2:**
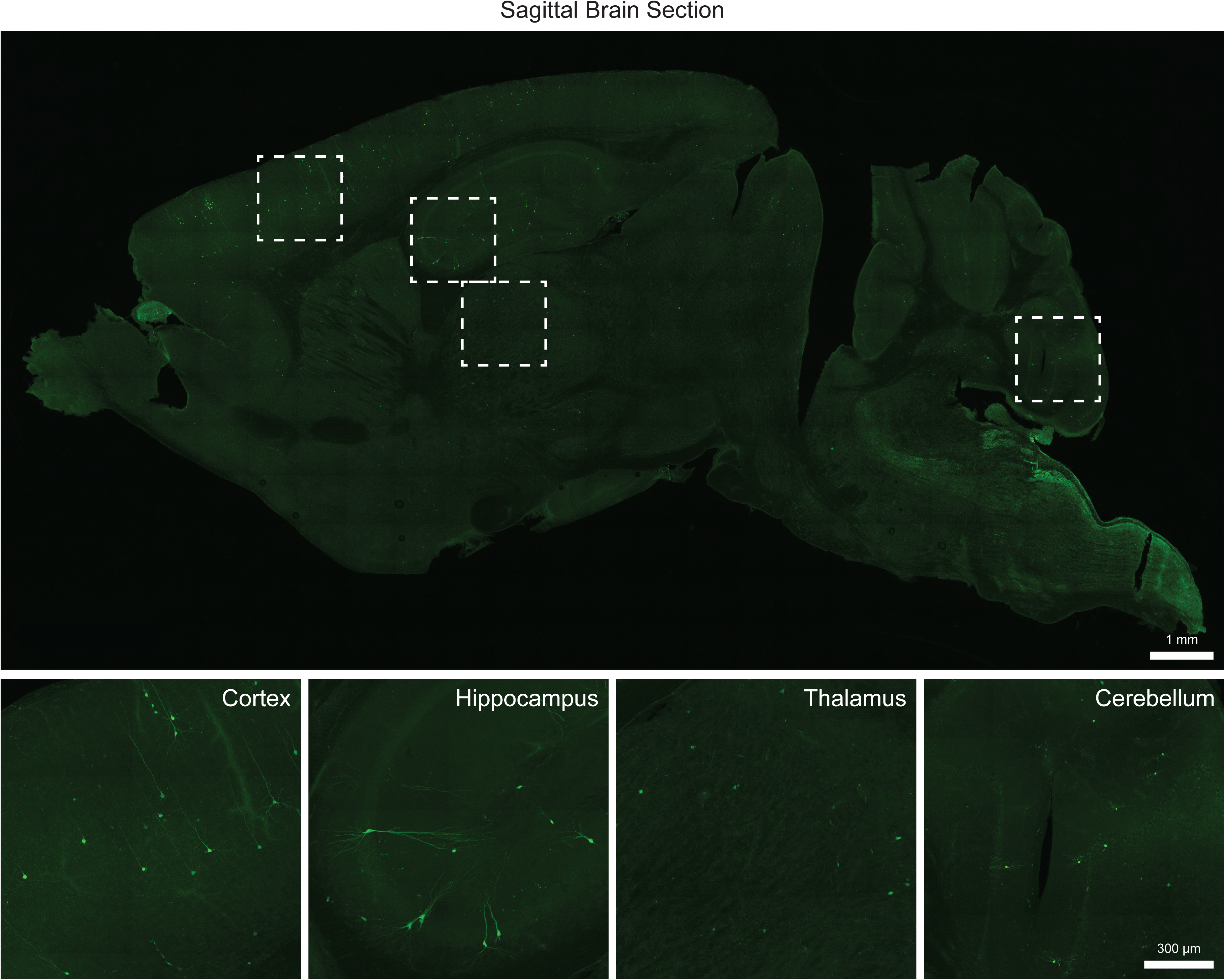
Characterization of MaCPNS1:hSyn-eGFP transduction in the *A. dimidiatus* brain. Top: Representative image of a mid-sagittal section of a brain stained for eGFP following systemic delivery of MaCPNS1:hSyn-eGFP (scale bar = 1 mm). Bottom: Enlarged images of the boxed regions showing representative areas of the cortex, hippocampus, thalamus, and cerebellum (scale bar = 300 μm).

## References

1. Allen, R. S. & Seifert, A. W. Spiny mice (Acomys) have evolved cellular features to support regenerative healing. Ann. N. Y. Acad. Sci. 1544, 5–26 (2025).

2. Okamura, D. M. et al. Spiny mice activate unique transcriptional programs after severe kidney injury regenerating organ function without fibrosis. iScience 24, 103269 (2021).

3. Seifert, A. W. et al. Skin shedding and tissue regeneration in African spiny mice (Acomys). Nature 489, 561–565 (2012).

4. Matias Santos, D., et al. Ear wound regeneration in the African spiny mouse Acomys cahirinus. Regeneration 3, 52–61 (2016).

5. Peng, H. et al. Adult spiny mice (Acomys) exhibit endogenous cardiac recovery in response to myocardial infarction. Npj Regen. Med. 6, 1–15 (2021).

6. Koopmans, T. et al. Ischemic tolerance and cardiac repair in the spiny mouse (Acomys). Npj Regen. Med. 6, 1–16 (2021).

7. Nogueira-Rodrigues, J. et al. Rewired glycosylation activity promotes scarless regeneration and functional recovery in spiny mice after complete spinal cord transection. Dev. Cell 57, 440–450.e7 (2022).

8. Streeter, K. A. et al. Molecular and histologic outcomes following spinal cord injury in spiny mice,. J. Comp. Neurol. 528, 1535–1547 (2020).

9. Thuret, S., Thallmair, M., Horky, L. L. & Gage, F. H. Enhanced Functional Recovery in MRL/MpJ Mice after Spinal Cord Dorsal Hemisection. PLOS ONE 7, e30904 (2012).

10. Dias, D. O. & Göritz, C. Fibrotic scarring following lesions to the central nervous system. Matrix Biol. 68-69, 561–570 (2018).

11. Stenimahitis, V. et al. Long-term outcome and predictors of neurological recovery in cervical spinal cord injury: a population-based cohort study. Sci. Rep. 14, 20945 (2024).

12. Brant, J. O., Lopez, M.-C., Baker, H. V., Barbazuk, W. B. & Maden, M. A Comparative Analysis of Gene Expression Profiles during Skin Regeneration in Mus and Acomys. PLOS ONE 10, e0142931 (2015).

13. Losner, J., Courtemanche, K. & Whited, J. L. A cross-species analysis of systemic mediators of repair and complex tissue regeneration. Npj Regen. Med. 6, 1–11 (2021).

14. Seifert, A. W., Duncan, E. M. & Zayas, R. M. Enduring questions in regenerative biology and the search for answers. *Commun*. Biol. 6, 1–13 (2023).

15. Gawriluk, T. R. et al. Comparative analysis of ear-hole closure identifies epimorphic regeneration as a discrete trait in mammals. Nat. Commun. 7, 11164 (2016).

16. Gawriluk, T. R. et al. Complex Tissue Regeneration in Mammals Is Associated With Reduced Inflammatory Cytokines and an Influx of T Cells. Front. Immunol. 11, (2020).

17. Tanaka, E. M. & Reddien, P. W. The Cellular Basis for Animal Regeneration. Dev. Cell 21, 172–185 (2011).

18. Simkin, J. et al. Tissue-resident macrophages specifically express *Lactotransferrin* and *Vegfc* during ear pinna regeneration in spiny mice. Dev. Cell 59, 496–516.e6 (2024).

19. Tomasso, A., Koopmans, T., Lijnzaad, P., Bartscherer, K. & Seifert, A. W. An ERK-dependent molecular switch antagonizes fibrosis and promotes regeneration in spiny mice (Acomys). Sci. Adv. 9, eadf2331 (2023).

20. Jozic, I., Stojadinovic, O., Kirsner, R. S. F. & Tomic-Canic, M. Skin under the (Spot)-Light: Cross-Talk with the Central Hypothalamic–Pituitary–Adrenal (HPA) Axis. J. Invest. Dermatol. 135, 1469–1471 (2015).

21. Lin, T.-K., Zhong, L. & Santiago, J. L. Association between Stress and the HPA Axis in the Atopic Dermatitis. Int. J. Mol. Sci. 18, 2131 (2017).

22. Marucha, P., Engeland, C. & Cacioppo, J. Wound Healing is Delayed in Women and in the Aged: A Potential Role for the HPA Axis. Wound Repair Regen. 12, A21–A21 (2004).

23. Elenkov, I. J., Wilder, R. L., Chrousos, G. P. & Vizi, E. S. The Sympathetic Nerve—An Integrative Interface between Two Supersystems: The Brain and the Immune System. Pharmacol. Rev. 52, 595–638 (2000).

24. Eijkelkamp, N., Engeland, C. G., Gajendrareddy, P. K. & Marucha, P. T. Restraint stress impairs early wound healing in mice via α-adrenergic but not β-adrenergic receptors. Brain. Behav. Immun. 21, 409–412 (2007).

25. Maydych, V. The Interplay Between Stress, Inflammation, and Emotional Attention: Relevance for Depression. Front. Neurosci. 13, (2019).

26. Saito, T., Tazawa, K., Yokoyama, Y. & Saito, M. Surgical stress inhibits the growth of fibroblasts through the elevation of plasma catecholamine and cortisol concentrations. Surg. Today 27, 627–631 (1997).

27. Alapure, B. V., Lu, Y., Peng, H. & Hong, S. Surgical Denervation of Specific Cutaneous Nerves Impedes Excisional Wound Healing of Small Animal Ear Pinnae. Mol. Neurobiol. 55, 1236–1243 (2018).

28. Skorvanek, M. & Bhatia, K. P. The Skin and Parkinson’s Disease: Review of Clinical, Diagnostic, and Therapeutic Issues. Mov. Disord. Clin. Pract. 4, 21–31 (2017).

29. Hui, Y. et al. Strategies for Targeting Neural Circuits: How to Manipulate Neurons Using Virus Vehicles. Front. Neural Circuits 16, (2022).

30. Haggerty, D. L., Grecco, G. G., Reeves, K. C. & Atwood, B. Adeno-Associated Viral Vectors in Neuroscience Research. Mol. Ther. - Methods Clin. Dev. 17, 69–82 (2020).

31. Boender, A. J. et al. An AAV-CRISPR/Cas9 strategy for gene editing across divergent rodent species: Targeting neural oxytocin receptors as a proof of concept. Sci. Adv. 9, eadf4950 (2023).

32. Wang, T., Teng, B., Yao, D. R., Gao, W. & Oka, Y. Organ-specific sympathetic innervation defines visceral functions. Nature 637, 895–902 (2025).

33. Mathon, B. et al. Increasing the effectiveness of intracerebral injections in adult and neonatal mice: a neurosurgical point of view. Neurosci. Bull. 31, 685–696 (2015).

34. Analysis of Transduction Efficiency, Tropism and Axonal Transport of AAV Serotypes 1, 2, 5, 6, 8 and 9 in the Mouse Brain | PLOS One. https://journals.plos.org/plosone/article?id=10.1371/journal.pone.0076310.

35. Watakabe, A. et al. Comparative analyses of adeno-associated viral vector serotypes 1, 2, 5, 8 and 9 in marmoset, mouse and macaque cerebral cortex. Neurosci. Res. 93, 144–157 (2015).

36. Gray, S. J. et al. Preclinical Differences of Intravascular AAV9 Delivery to Neurons and Glia: A Comparative Study of Adult Mice and Nonhuman Primates. Mol. Ther. 19, 1058–1069 (2011).

37. Yang, B. et al. Global CNS Transduction of Adult Mice by Intravenously Delivered rAAVrh.8 and rAAVrh.10 and Nonhuman Primates by rAAVrh.10. Mol. Ther. 22, 1299–1309 (2014).

38. Zincarelli, C., Soltys, S., Rengo, G. & Rabinowitz, J. E. Analysis of AAV Serotypes 1–9 Mediated Gene Expression and Tropism in Mice After Systemic Injection. Mol. Ther. 16, 1073–1080 (2008).

39. Majowicz, A. et al. Successful Repeated Hepatic Gene Delivery in Mice and Non-human Primates Achieved by Sequential Administration of AAV5ch and AAV1. Mol. Ther. 25, 1831–1842 (2017).

40. Halbert, C. L., Rutledge, E. A., Allen, J. M., Russell, D. W. & Miller, A. D. Repeat Transduction in the Mouse Lung by Using Adeno-Associated Virus Vectors with Different Serotypes. J. Virol. 74, 1524–1532 (2000).

41. Rivière, C., Danos, O. & Douar, A. M. Long-term expression and repeated administration of AAV type 1, 2 and 5 vectors in skeletal muscle of immunocompetent adult mice. Gene Ther. 13, 1300–1308 (2006).

42. Bedbrook, C. N., Deverman, B. E. & Gradinaru, V. Viral Strategies for Targeting the Central and Peripheral Nervous Systems. Annu. Rev. Neurosci. 41, 323–348 (2018).

43. Challis, R. C. et al. Adeno-Associated Virus Toolkit to Target Diverse Brain Cells. Annu. Rev. Neurosci. 45, 447–469 (2022).

44. Ravindra Kumar, S., et al. Multiplexed Cre-dependent selection yields systemic AAVs for targeting distinct brain cell types. Nat. Methods 17, 541–550 (2020).

45. Deverman, B. E. et al. Cre-dependent selection yields AAV variants for widespread gene transfer to the adult brain. Nat. Biotechnol. 34, 204–209 (2016).

46. Goertsen, D. et al. AAV capsid variants with brain-wide transgene expression and decreased liver targeting after intravenous delivery in mouse and marmoset. Nat. Neurosci. 25, 106–115 (2022).

47. Chen, X. et al. Functional gene delivery to and across brain vasculature of systemic AAVs with endothelial-specific tropism in rodents and broad tropism in primates. Nat. Commun. 14, 3345 (2023).

48. Chuapoco, M. R. et al. Adeno-associated viral vectors for functional intravenous gene transfer throughout the non-human primate brain. Nat. Nanotechnol. 18, 1241–1251 (2023).

49. Chen, X. et al. Engineered AAVs for non-invasive gene delivery to rodent and non-human primate nervous systems. Neuron 110, 2242–2257.e6 (2022).

50. Mathiesen, S. N., Lock, J. L., Schoderboeck, L., Abraham, W. C. & Hughes, S. M. CNS Transduction Benefits of AAV-PHP.eB over AAV9 Are Dependent on Administration Route and Mouse Strain. Mol. Ther. - Methods Clin. Dev. 19, 447–458 (2020).

51. Shay, T. F. et al. Primate-conserved carbonic anhydrase IV and murine-restricted LY6C1 enable blood-brain barrier crossing by engineered viral vectors. Sci. Adv. 9, eadg6618 (2023).

52. Huang, Q. et al. Delivering genes across the blood-brain barrier: LY6A, a novel cellular receptor for AAV-PHP.B capsids. PLOS ONE 14, e0225206 (2019).

53. Hordeaux, J. et al. The GPI-Linked Protein LY6A Drives AAV-PHP.B Transport across the Blood-Brain Barrier. Mol. Ther. 27, 912–921 (2019).

54. Jang, S., Shen, H. K., Ding, X., Miles, T. F. & Gradinaru, V. Structural basis of receptor usage by the engineered capsid AAV-PHP.eB. Mol. Ther. - Methods Clin. Dev. 26, 343–354 (2022).

55. Brittain, T. J. et al. Structural basis of liver de-targeting and neuronal tropism of CNS-targeted AAV capsids. 2025.06.02.655683 Preprint at 10.1101/2025.06.02.655683 (2025).

56. Quan, L., Uyeda, A. & Muramatsu, R. Central nervous system regeneration: the roles of glial cells in the potential molecular mechanism underlying remyelination. Inflamm. Regen. 42, 7 (2022).

57. Kim, Y. S., Choi, J. & Yoon, B.-E. Neuron-Glia Interactions in Neurodevelopmental Disorders. Cells 9, 2176 (2020).

58. Zhao, Y., Huang, Y., Cao, Y. & Yang, J. Astrocyte-Mediated Neuroinflammation in Neurological Conditions. Biomolecules 14, 1204 (2024).

59. Griffin, J. M. et al. Astrocyte-selective AAV gene therapy through the endogenous GFAP promoter results in robust transduction in the rat spinal cord following injury. Gene Ther. 26, 198–210 (2019).

60. Jagadisan, B. & Dhawan, A. Hepatotoxicity in Adeno-Associated Viral Vector Gene Therapy. Curr. Hepatol. Rep. 22, 276–290 (2023).

61. Griffiths, J. A. et al. Peripheral neuronal activation shapes the microbiome and alters gut physiology. Cell Rep. 43, (2024).

62. Wang, L., Yuan, P.-Q., Challis, C., Ravindra Kumar, S. & Taché, Y. Transduction of Systemically Administered Adeno-Associated Virus in the Colonic Enteric Nervous System and c-Kit Cells of Adult Mice. Front. Neuroanat. 16, (2022).

63. Cairns, B. R., Jevans, B., Chanpong, A., Moulding, D. & McCann, C. J. Automated computational analysis reveals structural changes in the enteric nervous system of nNOS deficient mice. Sci. Rep. 11, 17189 (2021).

64. Challis, R. C. et al. Systemic AAV vectors for widespread and targeted gene delivery in rodents. Nat. Protoc. 14, 379–414 (2019).

